# New data on Quaternary lagomorphs from Lagoa Santa, State Of Minas Gerais, Brazil

**DOI:** 10.1101/2025.11.10.687685

**Authors:** Victor Abade de Carvalho Ramos, Artur Chahud

## Abstract

Lagomorpha have a long evolutionary history, dating back to the Paleogene and currently exhibit a cosmopolitan distribution, inhabiting all continents except Antarctica. The present study analyzed and revised 148 fossil and subfossil specimens from Gruta Cuvieri, in Lagoa Santa (MG), attributed to the Late Pleistocene and Holocene. The methodology included sorting, anatomical identification of all skeletal elements, taxonomy, and taphonomy. The taxonomic analysis of a skull confirmed the presence of the species *Sylvilagus brasiliensis*, although it is not possible to rule out the occurrence of other species in the remaining assemblage, given the abundance of non-diagnostic bone elements. The taphonomy of the Cuvieri Cave revealed distinct patterns, with less reworked specimens in the Holocene deposits and greater dispersion in the Pleistocene deposits. The results indicate relevant differences in the fossil distribution and high fragmentation, suggesting distinct taphonomic processes associated with multiple depositional events.

## INTRODUCTION

The order Lagomorpha, which includes rabbits, hares, and pikas, is an important group of mammals that emerged during the Cenozoic. Fossil evidence indicates that lagomorphs originated in the Indian subcontinent during the Early Eocene, approximately 53 million years ago, based on small skeletal elements such as calcanei and astragali recovered from deposits in western India (Rose *et al*., 2008). In addition to this record, the known basal lagomorphs include *Dawsonolagus antiquus, Gomphos elkema*, and *Arnebolagus leporinus*, dated to the Early Eocene of Central Asia, China, and Mongolia (Li *et al*., 2007; Asher *et al*., 2005; Lopatin & Arianov, 2007).

The family Leporidae reached the North American continent during the Early Oligocene. The genus *Sylvilagus* Gray, 1867, which encompasses all South American species (Ruedas *et al*., 2017), originated in North America during the Late Pliocene to Early Pleistocene (Gazin, 1942; Emry & Gawne, 1986; White, 1991; Iraçabal *et al*., 2024).

In South America, the presence of lagomorphs is associated with the Great American Biotic Interchange (GABI), a phenomenon resulting from the formation of the Isthmus of Panama around 3 million years ago, which enabled faunal exchange between the Americas. It is estimated that the order entered South America approximately 125,000 years ago (Woodburne, 2010). The oldest records on the continent come from the Cangahua Formation in Ecuador (Ficarelli *et al*., 1992).

The Lagoa Santa region, in the state of Minas Gerais, hosts some of the most important archaeological and paleontological sites in Brazil, with a significant concentration of vertebrate fossils dated to the Late Pleistocene and Holocene (Alvarenga *et al*., 2008; Chahud, 2020a; Chahud, 2025; Chahud *et al*., 2021; Chahud & Okumura, 2021a; Chahud & Okumura, 2021b). Despite the relevance of these records, systematic studies comprehensively comparing Pleistocene and Holocene Leporidae fossils based on morphological, taxonomic, and ontogenetic criteria remain scarce.

The only studies on Leporidae from the Lagoa Santa region are those on fossils from the Cuvieri Cave, conducted by Chahud *et al*. (2020) and Chahud & Okumura (2022). This small cave contains a rich osteological assemblage, including a diverse Quaternary mammalian paleofauna such as tapirs (Chahud & Okumura, 2021b), armadillos (Chahud, 2021), pacas (Mayer *et al*., 2016; Chahud, 2022a), peccaries (Chahud & Okumura, 2023), and a large abundance of Cervidae remains (Chahud, 2020b; Chahud *et al*., 2024).

The present study aims to complement the works of Chahud *et al*. (2020) and Chahud & Okumura (2022) by incorporating data from newly identified specimens from Gruta Cuvieri, with emphasis on anatomical, taphonomic, and taxonomic aspects.

## MATERIALS AND METHODS

Cuvieri Cave is located within the Lagoa Santa Karst Environmental Protection Area (APA-Lagoa Santa Karts) and is named after the fossil of *Catonyx cuvieri* found at the site (Hubbe, 2008). Inside the cave, three small abysses occur, designated *Locus* 1, *Locus* 2, and *Locus* 3 (Figure 1), the latter subdivided into *Loci* 3A, 3B, and 3C.

**Figure 1.**
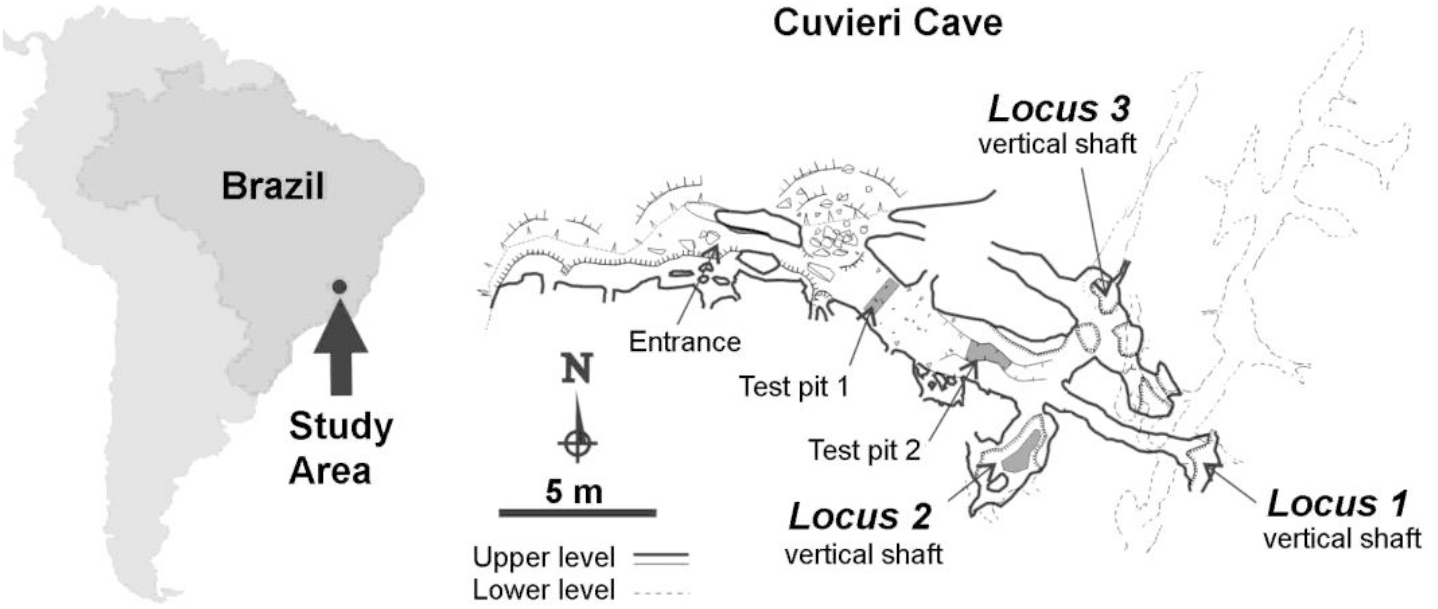
Geographic location of the study area and of Cuvieri Cave showing the position of *Loci* 1, 2 and 3. Map courtesy of Alex Hubbe and Grupo Bambuí for Speleological Research, 2020.

*Loci* 2 and 3 were subdivided into “exposures,” a term used to designate each excavation surface in which bones were exposed, documented, and subsequently recovered, functioning as proxies for the original depositional surfaces (Hubbe *et al*., 2011; Haddad-Martim *et al*., 2017). In total, *Locus* 2 was subdivided into 46 exposures; *Locus* 3A into 96 exposures; *Locus* 3B into 69 exposures; and *Locus* 3C into 4 exposures.

The *Loci* exhibit distinct depositional characteristics, with *Loci* 2 and 3 having been excavated with stratigraphic control. *Locus* 2 contains fossils dated to the entire Holocene and the Late Pleistocene, while *Locus* 3 predominantly preserves Pleistocene fossils (Hubbe, 2008).

The analysis was based on 148 fossils from Cuvieri Cave: 11 skeletal elements from *Locus* 1, 76 from *Locus* 2, and 61 from *Locus* 3 (59 from *Locus* 3A and only 2 from *Locus* 3B). Material selection considered criteria of integrity and morphological relevance, with emphasis on maxillae, mandibles, appendicular bones, and other diagnostic elements. Additionally, a skull from *Locus* 3A was examined to clarify its taxonomic classification.

Fossil screening considered their stratigraphic provenance, when available, and the feasibility of anatomical and taxonomic identification. Specimens assigned to Lagomorpha were separated from those of other small mammals, such as rodents, based on the identification of diagnostic morphological traits.

The selected fossils were subjected to descriptive morphological examination, documenting visible anatomical structures and preservation state (taphonomy). Specimens were catalogued in spreadsheets by anatomical element; ontogenetic stage (adult or subadult); modifications (such as concretion); and breakage level, classified according to the percentage of breakage, with 0% representing intact fossils and 80– 90% representing fragmented ones. A similar system was applied to concretion levels, with 0% indicating fossils without concretion and 80–90% indicating specimens with heavy concretion.

Postcranial anatomical and taxonomic identification was carried out through morphological comparison with a skeleton of *Oryctolagus cuniculus* (domestic rabbit). Although this species is not native to Brazil, its morphology is sufficiently similar to that of *Sylvilagus* to allow reliable comparisons, especially in the postcranium, where interspecific differences tend to be subtle. Anatomical and taxonomic analysis was complemented by specialized literature (Driesch, 1976; Paula Couto, 1979; Wible, 2007; Böhmer, 2015; Ruedas, 2017; Ruedas *et al*., 2017).

All analyzed specimens are registered and housed in the paleontological collection of the Laboratory for Human Evolutionary Studies at the Institute of Biosciences, University of São Paulo (IB-USP).

### SYSTEMATIC PALEONTOLOGY

Class Mammalia Linnaeus, 1758

Order Lagomorpha Brandt, 1855

Family Leporidae Fischer von Waldheim, 1817

Genus *Sylvilagus* Gray, 1867

*Sylvilagus brasiliensis* Linnaeus, 1758

Figures 2 and 3

**Figure 2.**
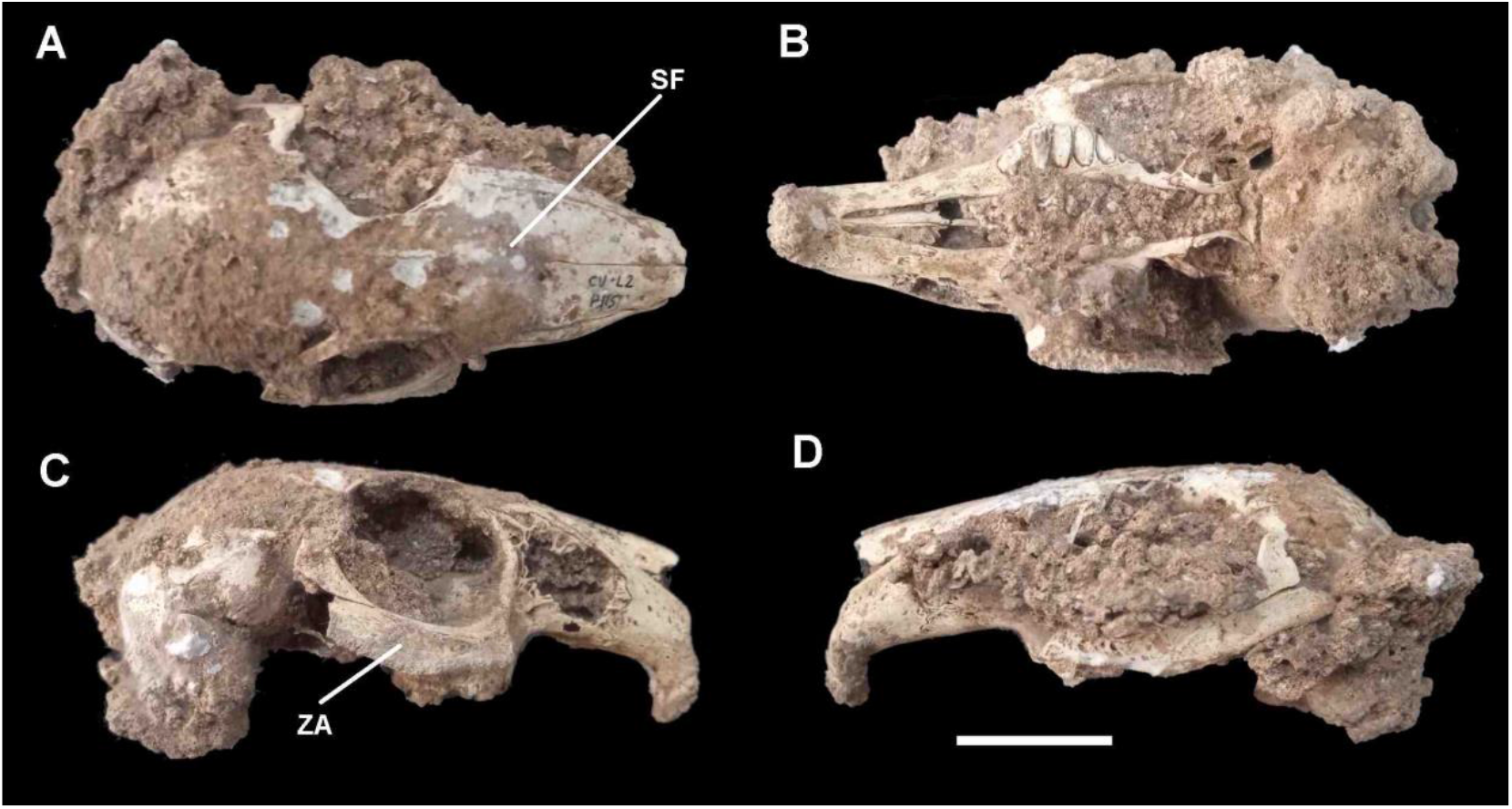
Skull of *Sylvilagus brasiliensis* from *Locus* 3A (CVL3-P115). A) Dorsal view; B) Ventral view; C and D) Lateral views. SF: suture frontonasal, ZA: Zygomatic Arch. Scale 10 mm.

**Figure 3.**
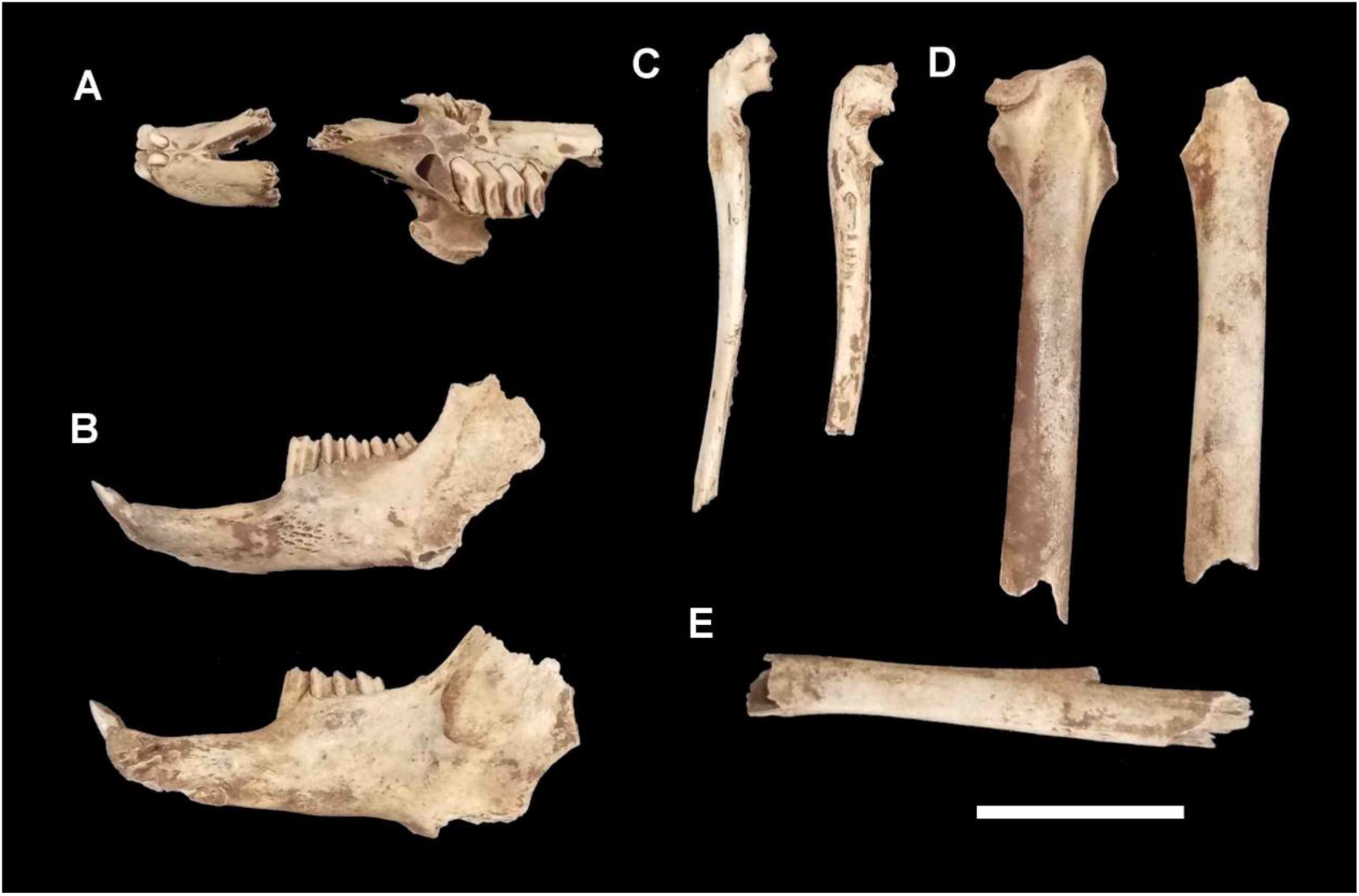
Leporidae specimen recovered from *Locus* 1. A) maxilla (CVL1-2b) and frontal portion (CVL1-2c); B) mandible split into two parts (CVL1-3a, CVL1-2a); C) right (CVL1-2d) and left (CVL1-2e) ulnae; D) right (CVL1-2f) and left (CVL1-2g) femora; fragmented tibia (CVL1-3b). Scale 20 mm.

#### Material

A fragmented and concretion-covered skull (Figure 2) from Exposure 1 of *Locus* 3A (CVL3-P115), along with several mandibles, maxillae, and other bone fragments from the three *loci*.

#### Remarks

The dentition, mandibular shape, and postcranial bones recovered from Cuvieri Cave are characteristic of the family Leporidae and can be attributed to the genus *Sylvilagus*, as no relevant morphological differences were observed among the analyzed skeletal elements.

Despite the high degree of concretion and the fragility of the specimen, it was possible to perform a taxonomic assessment of the skull (Figure 2), classifying it as *Sylvilagus brasiliensis*. This determination was based on the “U”-shaped frontonasal suture (Figure 2A), in contrast to the more “V”-shaped pattern observed in *Sylvilagus andinus* Thomas, 1897, and *Sylvilagus tapetillus* Thomas, 1913. Additional diagnostic traits include the long and closed postorbital process, distinct from the short or narrow process of *S. andinus* and *S. tapetillus*, and the strongly curved zygomatic arch (Figure 2C), which differs from the straighter condition described for the other species (Ruedas *et al*., 2017).

## Discussion

The skull CVL3-P115, from *Locus* 3A, exhibits diagnostic morphological characters consistent with *Sylvilagus brasiliensis*, confirming the presence of this species in the Lagoa Santa region during the Late Pleistocene. The mandibles described by Chahud *et al*.(2020) were found to be more elongated, slender, and gracile than those of *S. tapetillus*, and the mental foramen showed shape, size, and position compatible with *S. brasiliensis*.

Morphological comparisons performed by Chahud & Okumura (2022) demonstrated that the dentition and external morphology of the dentaries from Cuvieri Cave are indistinguishable from those of *Sylvilagus brasiliensis*. However, the mean mandibular series length (premolars + molars) was larger than that of the neotype of *S. brasiliensis* defined by Ruedas *et al*. (2017), as well as larger than the holotypes of *S. tapetillus* and *S. sanctaemartae* Hershkovitz, 1950. This larger body size pattern had already been noted by Lund and later by Paula Couto (1979), who referred to individuals from the region as *Sylvilagus brasiliensis minensis*. The validity of this subspecies, however, has been questioned, leading Ruedas *et al*. (2017) to propose raising *S. minensis* to species level for the larger specimens from southern and southeastern Brazil.

The postcranial bones recovered do not exhibit diagnostic traits allowing precise taxonomic identification. Thus, the presence of additional species within the fossil assemblage cannot be ruled out. It should be noted that no size differences were observed among adult specimens from the three *loci*.

Recent molecular studies (Silva *et al*., 2019) have revealed significant regional genetic differences among morphologically similar populations of *Sylvilagus*, reinforcing that isolated phenotypic variations are insufficient for species delimitation. In this context, the interpretation of fossil material must be carried out with caution, avoiding the attribution of size variation to taxonomic differences.

The analyzed material indicates the presence of both juvenile and adult individuals throughout the Pleistocene and Holocene layers, with no consistent morphological differences between them. Thus, the coexistence of multiple species cannot be confirmed based solely on morphological parameters, and the observed variation appears to reflect ontogenetic or population-level differences rather than specific divergence.

The identification of *Sylvilagus brasiliensis* is, therefore, the most appropriate for the skull specimen. The remaining skeletal elements, lacking diagnostic characters, are retained as *Sylvilagus* sp., as suggested by Chahud & Okumura (2022). Further comparative studies and genetic analyses involving fossils and subfossils of *Sylvilagus* from the Brazilian Quaternary are needed to elucidate their anatomical variability and historical distribution.

### TAPHONOMIC ANALYSIS

*Locus* 1 exhibited similar quantities of tibiae, femora, mandibles/maxillae, and ulnae, all located close to the surface, suggesting that they belonged to the same individual. It is likely that the specimen was relatively recent, with insufficient time for dispersal of its skeletal elements within the deposit (Figure 3).

The specimen shows dentition and dimensions consistent with an adult individual (Figures 3A and 3B). However, the disarticulated femoral epiphysis (Figure 3D) indicates that it was a subadult with a body size comparable to that of an adult.

This is the first study to present all osteological materials of Leporidae from *Locus* 1; previously, the specimnes had only been briefly mentioned by Chahud (2022b).

Previous studies conducted in other *loci* of Cuvieri Cave suggested a minimum number of six Lagomorpha individuals for *Locus* 3 and three for *Locus* 2 (Chahud; Mingatos; Okumura, 2020), calculated using the Minimum Number of Individuals (MNI) technique, which considers bone size and lateralization. However, taking into account the new materials and bone distribution, the actual number of individuals must have been significantly higher.

In *Loci* 2 and 3, it was possible to determine the exposure in which most fossils were recovered. In *Locus* 3, specimens are concentrated primarily in Exposures 1, 4, and 62 (Figure 5), whereas in *Locus* 2, Exposures 11 and 12 predominate (Figure 4).

**Figure 4.**
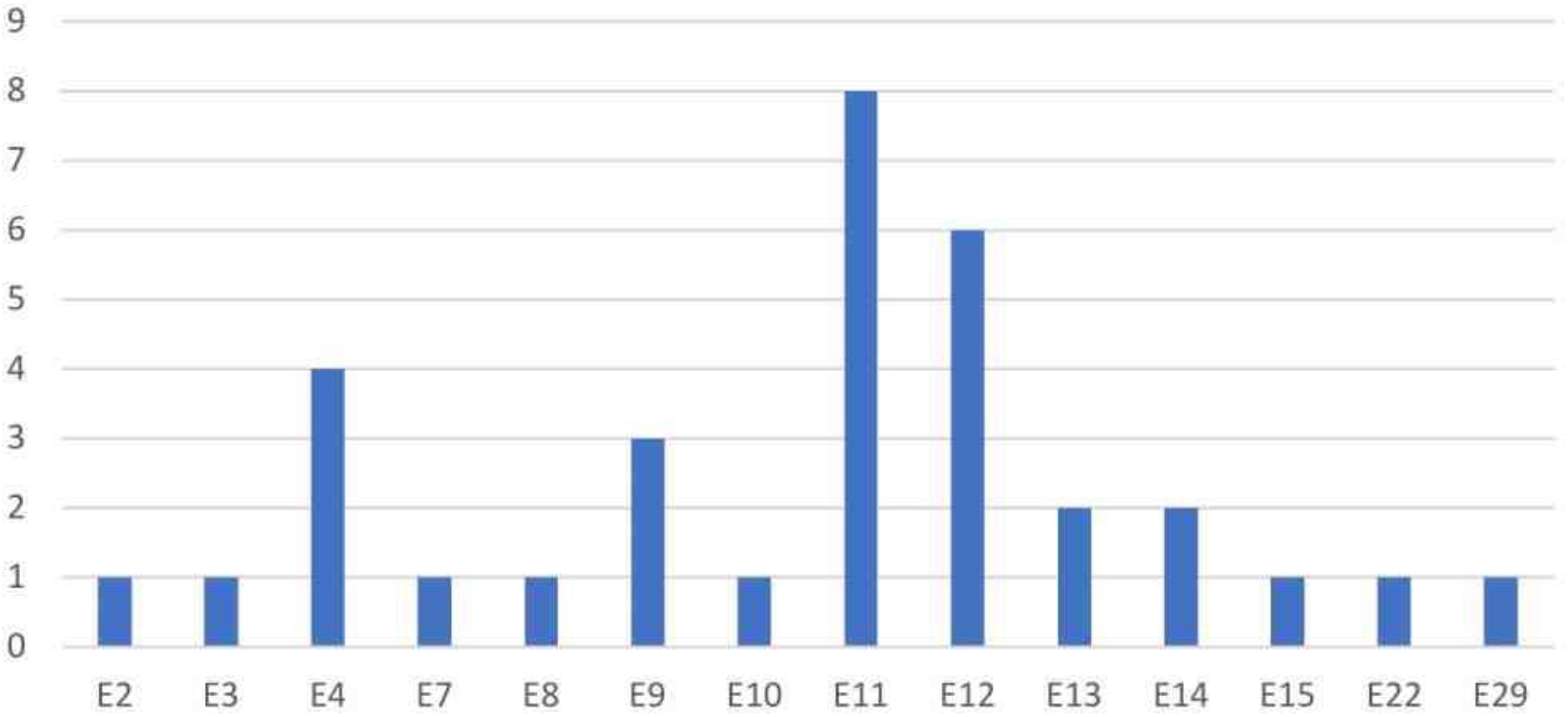
Number of pieces per exhibition in *Locus* 2

**Figure 5.**
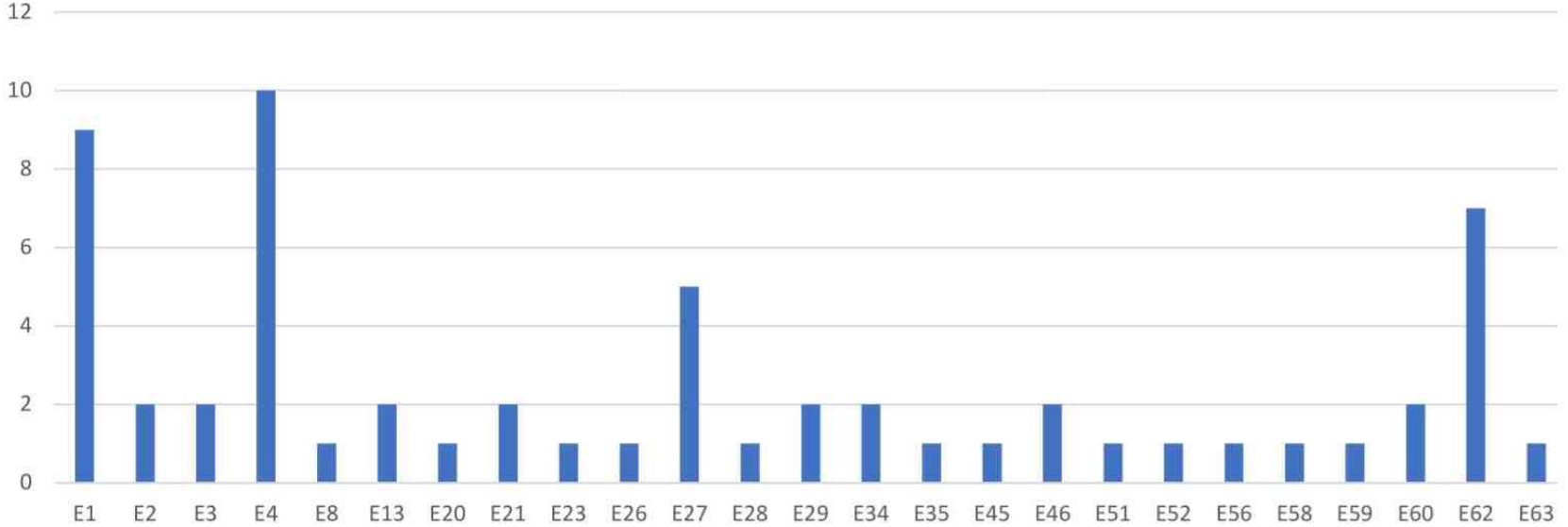
Number of pieces per display in *Locus* 3A

In the study by Chahud *et al*.(2020), no specimens were observed in exposures deeper than exposure 15, leading to the interpretation that all Leporidae from *Locus* 2 were younger than 6,000 years, since specimens found between exposures 11 and 17 were estimated to be of that age (Hubbe *et al*., 2011). However, the discovery of two maxillae in exposures 22 and 29 suggests that the genus *Sylvilagus* was present in the Lagoa Santa region earlier than 7,000 years ago, throughout much of the Middle Holocene, although it had not yet become as common as observed in younger deposits.

The distribution of specimens across various exposures in *Locus* 3 indicates reworking processes but confirms the presence of *Sylvilagus* during the Pleistocene. Recently, Fabris *et al*. (2025) identified a specimen older than 9,000 years at the Lapa do Santo archaeological site, also in the Lagoa Santa region, demonstrating that the genus was already present at the beginning of the Holocene.

A significant difference in anatomical element distribution was observed between *loci* (Table 1), with *Locus* 3 yielding a larger number of tibiae compared to other bones, whereas *Locus* 2 contained more molars and incisors. More fragile elements, such as vertebrae, were also recorded in *Locus* 2, suggesting reduced reworking and remobilization compared to *Locus* 3, which favored the preservation of delicate structures.

**Table 1.** Number of skeletal elements in *Loci* 2 and 3.

|  | *Locus 2* | *Locus 3* |
| --- | --- | --- |
| Skull | 0 | 1 |
| Maxilla fragments | 7 | 2 |
| Hemimandibles | 5 | 12 |
| Isolated incisors | 11 | 1 |
| Isolated premolars | 1 | 0 |
| Isolated molars | 15 | 0 |
| Scapula | 1 | 6 |
| Humerus | 3 | 5 |
| Radio | 1 | 7 |
| Ulna | 1 | 0 |
| Vertebra | 11 | 3 |
| Sacrum | 0 | 1 |
| Rib | 1 | 0 |
| Femur | 3 | 10 |
| Tibia | 3 | 6 |
| Pelvic bones | 3 | 5 |
| Calcaneus | 5 | 0 |
| Metatarsus | 1 | 0 |
| Indeterminate Metapodium | 3 | 0 |
| Phalanx | 1 | 0 |
| Indeterminate | 0 | 2 |
| **TOTAL** | **76** | **60** |

*Locus* 3C did not contain specimens, while *Locus* 3B yielded only two scapular fragments, indicating that leporids were predominantly restricted to *Locus* 3A, which is closer to the main conduit of Cuvieri Cave.

High fragmentation levels, ranging from intact fossils (0%) to specimens with more than 90% breakage (Figure 6), indicate the action of intense remobilization processes within the cave. This remobilization may have a biological origin, such as trampling or the fall of large animals into the cavity, or may result from sediment movement caused by water flow.

**Figure 6.**
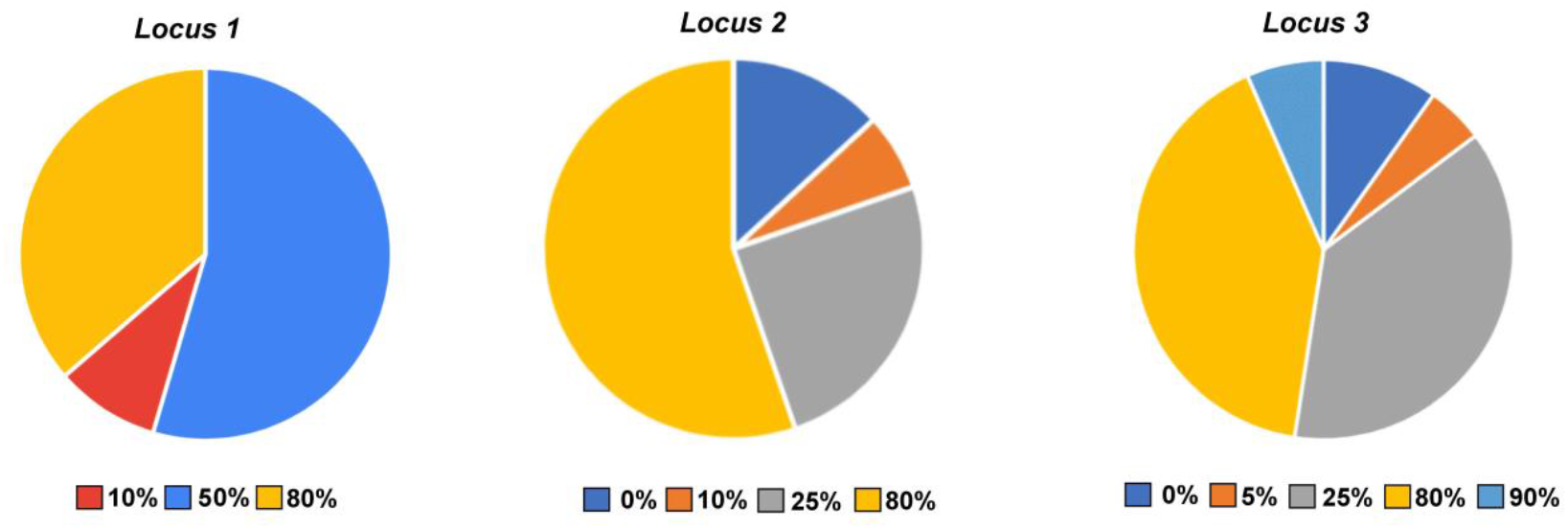
Percentage of breakage of parts from *Loci* 1, 2, and 3

The distinction between exposures and the anatomical distribution of fossils among the *loci* reinforces the hypothesis of different preservation conditions over time. Differences in anatomical representation, such as the predominance of teeth and a greater number of fragile bones (vertebrae) in *Locus* 2, and the more frequent presence of hemimandibles and long bones in *Locus* 3, suggest bone selection patterns similar to those described for medium-sized mammals reworked by hydraulic flow (Voorhies, 1969).

Although hydraulic sorting likely occurred, it was probably not intense, considering that articulated remains of medium and large sized mammals were found in all *loci*. Biological activity, especially the fall of large animals, is more likely to have been the main agent responsible for bone fragmentation and selective preservation. This interpretation aligns with the pattern observed in *Locus* 3, where remains of large-bodied taxa, such as ground sloths, adult tapirs, cervids, large tayassuids, and large rodents, are common (Hubbe, 2008; Mayer *et al*., 2016; Chahud *et al*., 2023a; 2023b) and could have destroyed smaller bones and displaced larger, more resistant elements. This hypothesis is reinforced by the greater number of specimens found both in shallow exposures and in the deeper exposures of *Locus* 3, where large mammals were not recorded.

In *Locus* 2, the largest mammals present in levels containing Leporidae are Cervidae and the tayassuid *Dicotyles tajacu* (Chahud & Okumura, 2023), and their destructive impact was likely lower than in *Locus* 3, although still significant.

In contrast to the high fragmentation levels, the low concretion percentages (Figure 7), rarely exceeding 80% even in the most altered specimens, indicate relatively homogeneous depositional conditions, with no mineral cementation processes capable of heavily coating bone surfaces. This is relevant because it supports the hypothesis of relatively rapid burial and limited exposure for most specimens. Although some fossils exhibit concretions that hinder anatomical analysis, as in the case of the described skull, most specimens remained relatively unaffected by this type of alteration.

**Figure 7.**
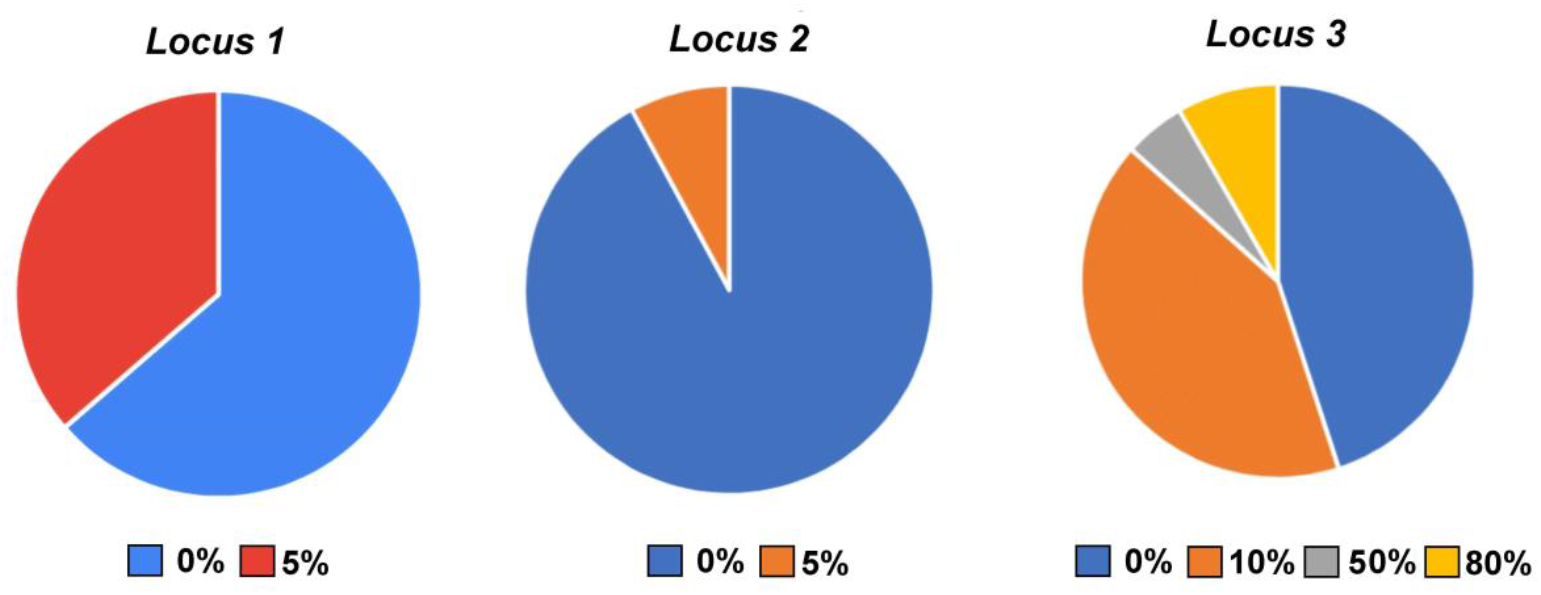
Percentage of concrete formation of the pieces from *Loci* 1, 2, and 3

## CONCLUSIONS

The analysis of fossils attributed to the genus *Sylvilagus* from Cuvieri Cave, in Lagoa Santa (MG), enabled the identification of *Sylvilagus brasiliensis* based on a skull recovered from *Locus* 3. However, specimens represented only by postcranial elements could not be identified at the species level and remain classified as *Sylvilagus* sp., as proposed by Chahud & Okumura (2022). No significant morphological differences were observed among specimens from the different *loci*.

The new specimens found in *Locus* 2 confirm the presence of *Sylvilagus* in Middle Holocene deposits, although its occurrence during this period appears to be rare compared to the Pleistocene and Late Holocene. In the Pleistocene, represented by *Locus* 3, the family Leporidae was present and was likely common in the late part of the period.

Deposition in *Loci* 2 and 3 differed, with the Pleistocene material from *Locus* 3 being more reworked and remobilized than the Holocene material from *Locus* 2. The low concretion levels observed across all loci suggest relatively brief exposure of specimens, whereas the high fragmentation levels indicate that biological agents, particularly the fall of large animals, were the primary taphonomic factors. This process is most evident in *Locus* 3, where the highest number of specimens occurs in exposures with few or no associated large mammals.

